# Triticeae resources in Ensembl Plants

**DOI:** 10.1101/011585

**Authors:** Dan M. Bolser, Arnaud Kerhornou, Brandon Walts, Paul Kersey

## Abstract

Recent developments in DNA sequencing have enabled the large and complex genomes of many crop species to be determined for the first time, even those previously intractable due to their polyploid nature. Indeed, over the course of the last two years, the genome sequences of several commercially important cereals, notably barley and bread wheat, have become available, as well as those of related wild species. While still incomplete, comparison to other, more completely assembled species suggests that coverage of genic regions is likely to be high.

Ensembl Plants (http://plants.ensembl.org) is an integrative resource organising, analysing and visualising genome-scale information for important crop and model plants. Available data includes reference genome sequence, variant loci, gene models and functional annotation. For variant loci, individual and population genotypes, linkage information and, where available, phenotypic information, are shown. Comparative analyses are performed on DNA and protein sequence alignments. The resulting genome alignments and gene trees, representing the implied evolutionary history the gene family, are made available for visualisation and analysis. Driven by the use case of bread wheat, specific extensions to the analysis pipelines and web interface have recently been developed to support polyploid genomes.

Data in Ensembl Plants is accessible through a genome browser incorporating various specialist interfaces for different data types, and through a variety of additional methods for programmatic access and data mining. These interfaces are consistent with those offered through the Ensembl interface for the genomes of non-plant species, including those of plant pathogens, pests and pollinators, facilitating the study of the plant in its environment.

## Introduction

The first cereal genome sequenced was rice in 2002 (Goff et al., 2002; Yu et al., 2002). More recently, progress has accelerated with the publication of the genome sequence of maize in 2009 (Schnable et al., 2009), barley in 2012 (IBSC, 2012), progenitors of the bread wheat A and D genomes in 2013 (Jia et al., 2013; Ling et al., 2013), and the draft bread wheat genome itself in 2014 (IWGSC, 2014; Brenchley et al., 2012). These four cereals, barley, maize, rice and wheat, together account for 30% of global food production or 2.4 out of 3.8 billion tonnes annually.

It is important to note that the current reference genome assemblies vary considerably in their contiguity and in the detail of available functional annotation. The Triticeae genomes were all sequenced primarily using short read sequencing (mainly from the Illumina platform), and the completion of these assemblies remains a scientific challenge, due to their large size and repetitive natures. The improvement of sequencing technologies, particularly those capable of capturing long-range information, will be necessary to achieve this goal. However, even in their existing condition, these resources are already sufficiently complete to be usefully represented through data analysis and visualisation platforms designed for genomes with finished assemblies, such as Ensembl Plants.

Ensembl Plants (http://plants.ensembl.org) offers integrative access to a wide range of genome-scale data from plant species (Kersey et al., 2014), using the Ensembl software infrastructure (Flicek et al., 2014). Currently, the site includes data from 38 plant genomes, from algae to flowering plants. Genomes are selected for inclusion in the resource based on the availability of complete genome sequence, their importance as model organisms (e.g. **Arabidopsis thaliana**, **Brachypodium distachyon**), their importance in agriculture (e.g. potato, tomato, various cereals and Brassicaceae), or because of their interest as evolutionary reference points (e.g the basal angiosperm, *Amborella trichopoda*, the aquatic alga *Chlamydomonas reinhardtii*, the moss *Physcomitrella patens* and the vascular non-seed spikemoss *Selaginella moellendorffii*). In total, the resource contains the genomes of 19 true grasses, *Musa accuminata* (banana), 12 dicots and 6 other species that provide evolutionary context for the plant lineage.

All species in the resource have data for genome sequence, annotations of protein-coding and non-coding genes and gene-centric comparative analysis. Additional data types within the resource include gene expression, sequence polymorphism, and whole genome alignments, which are selectively available for different species. In this sense, Ensembl Plants is similar to comparable, species, clade or data-type specific resources such as WheatGenome.info (Lai et al., 2012), HapRice (Yonemaru et al., 2014) or ATTED-II (Obayashi et al., 2014).

Ensembl Plants is released 4-5 times a year, in synchrony with releases of other genomes (from animals, fungi, protists and bacteria) in the Ensembl system. The provision of common interfaces allows access to genomic data from across the tree of life in a consistent manner, including data from plant pathogens, pests and pollinators.

## Database

### The Ensembl genome browser

Interactive access to Ensembl Plants is provided through an advanced genome browser. The browser allows users to visualise a graphical representation a completely assembled chromosome sequence or a contiguous sequence assembly comprising only a small portion of a molecule at various levels of resolution. Functionally interesting “features” are depicted on the sequence with defined locations. Features include conceptual annotations such as genes and variant loci, sequence patterns such as repeats, and experimental data such as sequence features mapped onto the genome, which often provide direct support for the annotations (Figure 1). Functional information is provided through import of manual annotation from the UniProt Knowledgebase (The Uniprot Consortium, 2014), imputation from protein sequence using the classification tool InterProScan (Jones et al., 2014), or by projection from orthologues (described below). Users can download much of the data available on each page in a variety of formats, and tools exist for upload of various types of user data, allowing users to see their own annotation in the context of the reference sequence. DNA and protein-based sequence search are also available. All genomes included in Ensembl Plants are periodically run through the Ensembl comparative analysis pipelines, generating DNA and protein sequence alignments. Gene trees are based on protein sequence alignments and show the inferred evolutionary history of each gene family (Vilella et al., 2009). Specialised views are available for these data (for example, see Figure 2, 5 and 6), and also for data types including variation (Figure 3), regulation, and expression.

**Figure 1:**
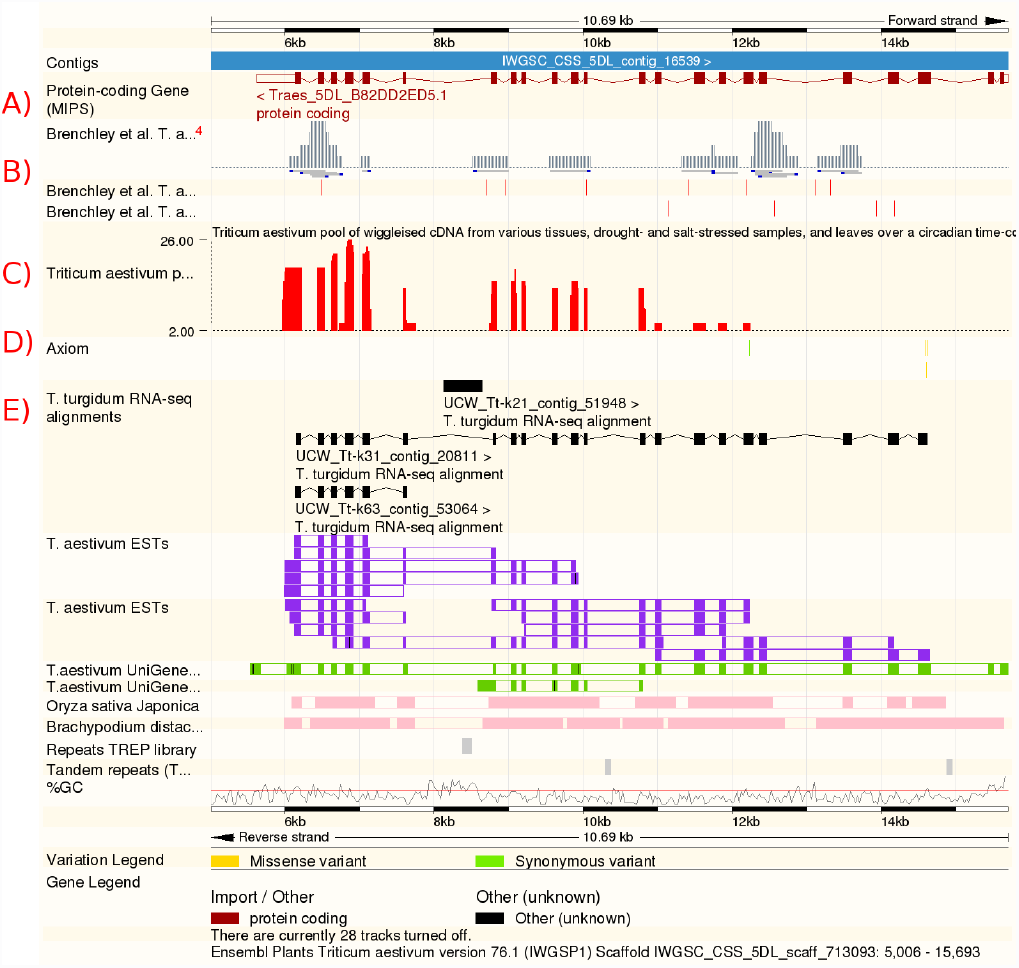

Visualising the bread wheat genome through the Ensembl Genomes browser interface. The user can view many layers of genome annotation in a highly customisable way. Tracks shown include: A) gene models, B) Assemblies and inter-homoeologous variations from Brenchley et al., C) RNA-Seq data, D) variations from the AXIOM array, and E) transcript assemblies from *T. turgidum*. Additional tracks are shown for *T. aestivum* ESTs and UniGenes (purple and green), alignment blocks to *O. sativa* and *B. distachyon* (pink), repeats (grey) and GC content.

**Figure 2:**
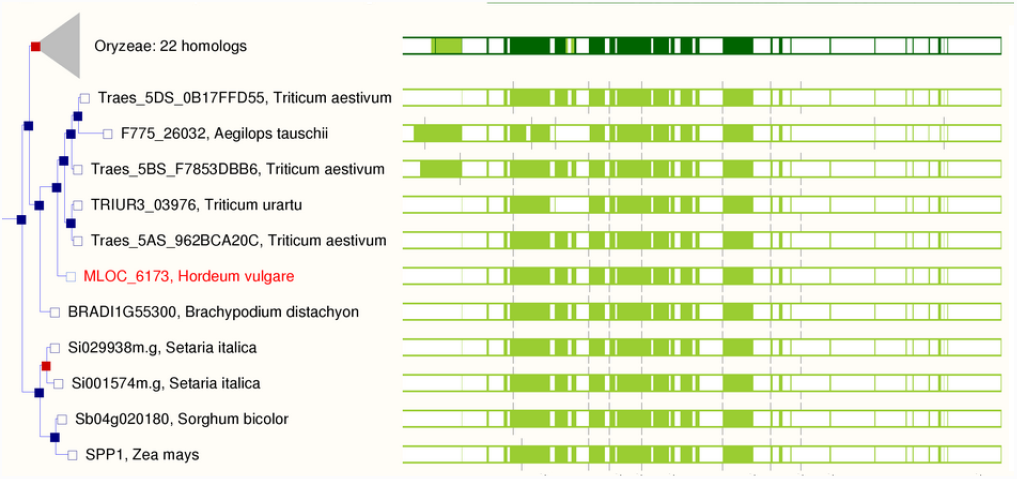

Detailed view of a gene tree in Ensembl Plants. The tree shows the inferred evolutionary history of the Sucrose-6F-phosphate phosphohydrolase family protein in *H. vulgare*. The gene tree (left) shows the expected phylogenetic relationship for the gene between the species shown. Note that the sequence identifier of the wheat genes includes the name of the chromosome arm to which it belongs, i.e. 5DS for the short arm of chromosome 5 in the D-genome. Red squares indicate inferred duplication events in the history of the gene, and shaded grey triangles indicate collapsed branches. A pictographic representation of the underlying multiple sequence alignment is included on the gene tree pages (right).

**Figure 3:**
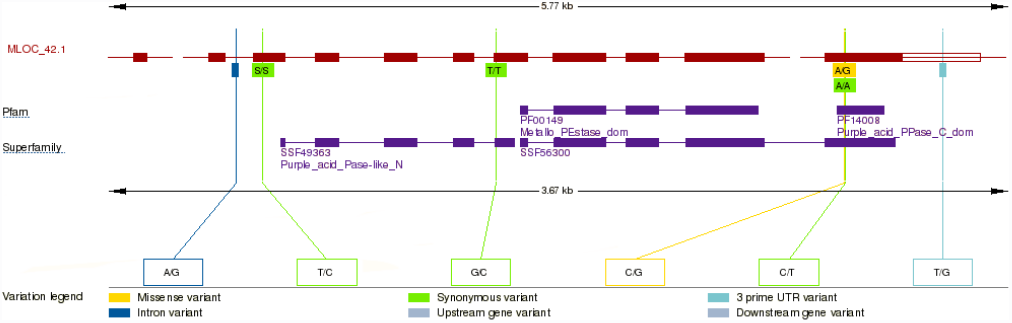

The transcript variation image for the *H. vulgare* MLOC_42.1 protein-coding transcript in Ensembl Plants. The image gives an overview of all the variants within the transcript in the context of the functional domains assigned to the protein. Upper boxes highlight the amino acid change, where applicable, and lower boxes give the alleles. Variants are colour coded according to their consequence type, missense, synonymous, and positional. A full list of consequence types is given here: http://www.ensembl.org/info/genome/variation/predicted_data.html. The transcripts, features and variations can be clicked to explore more information about each object.

The data are stored in MySQL databases using the same schemas as those used by other Ensembl sites. Direct access to these is provided through a public MySQL server and additionally through well-developed Perl and REST APIs. Database dumps and common data sets, such as DNA, RNA, protein sequence sets and sequence alignments, can be directly downloaded in bulk via FTP (ftp://ftp.ensemblgenomes.org).

In addition to the primary databases, Ensembl Plants also provides access to denormalised data warehouses, constructed using the BioMart toolkit (Kasprzyk, 2011). These are specialised databases optimised to support the efficient performance of common gene- and variant-centric queries, and can be accessed through their own web-based and programmatic interfaces.

### Triticeae genomes in Ensembl Plants

Four Triticeae genomes are currently hosted in Ensembl Plants (Table 1): *Hordeum vulgare* (barley), *Triticum aestivum* (bread wheat, also known as common wheat), and the genomes of two of bread wheat’s diploid progenitors: *Triticum urartu* (the A-genome progenitor) and *Aegilops tauschii* (the D-genome progenitor). In addition, a further three wheat transcriptomes were included by alignment (described below).

**Table 1:** Triticeae genomes in Ensembl Plants.

| Species (strain) | Description | Estimated genome size Mb | Assembly size Mb | Genes |
| --- | --- | --- | --- | --- |
| Barley<br>*Hordeum vulgare*<br>(cv. Morex) | Barley is an economically important crop and an important model of environmental diversity for development of wheat (IBSC, 2012). | 5,100<br>(DOLEŽEL et al., 1998) | 4,706 | 24,211 |
| Bread wheat<br>*Triticum aestivum*<br>(cv. Chinese spring) | An economically important food crop, accounting for over 20% of global agricultural production (IWGSC, 2014). | 16,974<br>(Bennett and Smith, 1991) | 4,460 | 108,569 |
| A-genome progenitor<br>*Triticum urartu*<br>(G1812) | An einkorn wheat and the diploid progenitor of the bread wheat A-genome (Ling et al., 2013). | 4,940<br>(Ling et al., 2013) | 3,747 | 34,843 |
| D-genome progenitor<br>*Aegilops tauschii*<br>(AL8/78) | The diploid progenitor of the bread wheat D-genome (Jia et al., 2013). | 4,360<br>(Jia et al., 2013) | 3,314 | 33,849 |

Barley is the world's fourth most important cereal crop and an important model for ecological adaptation, having been cultivated in all temperate regions from the Arctic Circle to the tropics. It was one of the first domesticated cereal grains, originating in the Fertile Crescent of south-west Asia / north-east Africa over 10,000 years ago (Harlan and Zohary, 1966). With a haploid genome size of approximately 5.3 Gbp in 7 chromosomes, the barley genome is among the largest yet sequenced. However, as a diploid, it is a natural model for the genetics and genomics of the polyploid members of Triticeae tribe, including wheat and rye.

The current barley genome assembly (cv. Morex) was produced by the International Barley Genome Sequencing Consortium (IBSC, 2012). The assembly is highly fragmented, but comparison to related grass species suggested that coverage of the gene space was good. The assembly was dubbed a ‘gene-ome’, a near complete gene set integrated into a chromosome-scale assembly using physical and genetic information. Sequence contigs that could not be assigned chromosomal positions in this way, were binned by homology to low-coverage shotgun sequence of flow-sorted chromosome arms (Muñoz-Amatriaín et al., 2011). This method integrated 22% of the total assembled sequence by length, covering 91% of the genes, into the chromosome-scale assembly.

Bread wheat is a major global cereal grain, essential to human nutrition. The bread wheat genome is hexaploid, with a size estimated at ~17 Gbp, composed of three closely related and independently maintained genomes (the A, B and D genomes). This complex structure has resulted from two independent hybridisation events. The first event brought together the diploid *T. urartu* (the A-genome donor) and an unknown Aegilops species, thought to be related to *Aegilops speltoides* (the B-genome donor), forming the allotetraploid *T. turgidum* around 0.5 MYA. This species has produced both the emmer and durum wheat cultivars, the latter being still grown today for pasta. The second hybridisation event brought together *T. turgidum* with *Ae. tauschii* (the D-genome donor) about 8,000 years ago in the Fertile Crescent.

Ensembl Plants contains version 1.0 of the chromosome survey sequence for *T. aestivum* cv. Chinese Spring, generated by the International Wheat Genome Sequencing Consortium (IWGSC, 2014). The draft assembly of this complex genome was made tractable by using flow-sorted chromosome arms (Dolezel et al., 2007; Vrána et al., 2012). The assembly of gene-containing regions is reasonably good, with an N50 of 2.5kb (Table 2).

**Table 2:** Some gene model statistics. Contig N50 reported twice, once for the complete assembly / and once for just the gene containing contigs. Complete genes are defined as those starting with a methionine, ending in a stop codon. InterPro coverage consists only of structural protein domains and functional motifs, excluding low complexity, coiled coil, transmembrane and signal motifs

| Species | Contig N50<br>(kbp) | Average<br>gene length | Average<br>exon number | % complete<br>genes | % InterPro<br>coverage |
| --- | --- | --- | --- | --- | --- |
| *O. sativa* |  | 3,090 | 2.5 | 77 | 68 |
| *B. distachyon* |  | 3,582 | 5.6 | 99 | 86 |
| *H. vulgare* | 0.9 / 3.2 | 2,812 | 5.4 | 76 | 84 |
| *T. urartu* | 3.4 / 5.8 | 3,208 | 4.7 | 78 | 78 |
| *Ae. tauschii* | 4.5 / 6.2 | 2,935 | 4.9 | 100 | 77 |
| *T. aestivum* | 2.4 / 6.3 | 2,197 | 3.8 | 56 | 84 |

The draft genome assemblies of *Ae. tauschii* (AL8/78) and *T. urartu* (G1812) are 4.23 and 4.66 Gbp, with N50 lengths of 58 kbp and 64 kbp, respectively. They were both produced by the Chinese Academy of Agricultural Sciences using similar methodologies (Jia et al., 2013; Ling et al., 2013)

## Data import

Gene model annotations and gene names were imported from the relevant authority for each species (see references in Table 1). For specific genomes, additional sequence and variation datasets have been added, and are described below for the four Triticeae genomes in Ensembl Plants. Information about the analysis and visualisation of the available data is described in the Section “Analysis and visualisation”.

### H. vulgare (barley)

A total of 79,379 gene models were described with the release of the barley genome (IBSC, 2012). These models are classified as either high-confidence (26,159 genes) or low-confidence (53,220 genes), which are displayed in separate tracks in the browser. Only the high-confidence gene models were used for downstream analysis (see “Analysis and visualisation”, below). The gene models for barley were made available through the MIPS barley genome database (http://mips.helmholtz-muenchen.de/plant/barley/).

Expressed sequences including Expressed Sequence Tags (ESTs) from the HarvEST database (http://harvest.ucr.edu) and a set of non-redundant barley full-length cDNAs (Matsumoto et al., 2011) were aligned to the genome to demonstrate support for the gene models. Sequences from the Affymetrix GeneChip Barley Genome Array (http://www.affymetrix.com/catalog/131420/AFFY/Barley-Genome-Array) were also aligned, allowing users to search the genome by probe identifier and find the corresponding regions or transcripts in barley.

RNA-Seq data was aligned from two studies: (i) SNP Discovery in Nine Lines of Cultivated Barley (Study ERP001573) and (ii) RNA-Seq study of eight growth stages (Study ERP001600), both described in the reference publication (IBSC, 2012).

Sequence variation was loaded from two sources: 1) variants derived from the whole genome shotgun survey sequencing four cultivars, Barke, Bowman, Igri, Haruna Nijo and a wild barley, *H. spontaneum*, and 2) variants derived from RNA-Seq from the embryonic tissues of nine spring barley varieties (Barke, Betzes, Bowman, Derkado, Intro, Optic, Quench, Sergeant and Tocada). Both approaches are described in the reference publication (IBSC, 2012). Figure 3 shows an example view of barley variations in Ensembl Plants. See below for more information on analysis of variation data.

### T. aestivum (bread wheat)

A total of 99,386 gene models were described with the release of the wheat chromosome survey sequence (IWGSC, 2014). Their structure was computed by spliced-alignments of publically available wheat full-length cDNAs and the protein sequences of the related grass species, barley, brachypodium, rice and Sorghum. A comprehensive RNA-seq dataset including five different tissues, root, leaf, spike, stem and grain, and different developmental stages was also used to identify wheat specific genes and splice variants (Figure 1A). The gene models for wheat were made available through the MIPS Wheat Genome Database (http://mips.helmholtz-muenchen.de/plant/wheat/).

A set of wheat genome assemblies (Brenchley et al., 2012) were aligned to the IWGSC assembly as well as to brachypodium, barley and the wheat progenitor genomes. Homoeologous variants that were inferred between the three component wheat genomes in the same study are also displayed in Ensembl Plants in the context of the gene models of these five species (Figure 1B).

RNA-Seq data was aligned from three studies: (i) Discovery of SNPs and genome-specific mutations by comparative analysis of transcriptomes of hexaploid wheat and its diploid ancestors (Study SRP002455; Akhunova et al., 2010) (ii) 454 sequencing of the *T. aestivum* cv. Chinese spring transcriptome (Study ERP001415; Brenchley et al., 2012), and (iii) *T. aestivum* Transcriptome or Gene expression (Study SRP004502) (Figure 1C).

Variations for bread wheat were loaded from CerealsDB (Wilkinson et al., 2012). A total of approximately 725,000 SNPs across approximately 250 varieties were loaded. These SNP loci are associated with markers from three genotyping platforms: The Axiom 820K SNP Array, the iSelect 80K Array (Wang et al., 2014), and the KASP probeset (Allen et al., 2011). In addition to these inter-varietal SNPs, work is ongoing to generate and report inter-homoeologous variants (Figure 1D).

### T. urartu and Ae. tauschii (the bread wheat A and D progenitor genomes)

A total of 34,879 and 34,498 protein-coding genes were reported for *T. urartu* and *A tauschii*, respectively. They were predicted using FGENESH (Salamov and Solovyev, 2000) and GeneID (Guigó et al., 1992) with supplemental evidence from RNA-Seq and EST sequences (Jia et al., 2013; Ling et al., 2013). In addition, approximately 200,000 bread wheat UniGene cluster sequences were aligned to both genomes using Exonerate (Slater and Birney, 2005). UniGene cluster sequences are based on reads from cDNA and EST libraries across a variety of samples (Wheeler et al., 2003). Similar sequences are clustered into transcripts, and in this case, are filtered by species (*T. aestivum*). Similarly, all publicly available bread wheat ESTs, retrieved using the European Nucleotide Archive (Leinonen et al., 2011) advanced search, were aligned using STAR (Dobin et al., 2013).

For *Ae. tauschii*, RNA-Seq data was aligned from a single study: RNA-Seq from seedling leaves of *Ae. tauschii* (Study DRP000562; Iehisa et al., 2012).

### Transcriptomes

In addition to the four Triticeae genomes, the transcriptome assembly of *Triticum turgidum* (durum wheat) is presented by alignment to *T. aestivum* and the transcriptome assemblies of two *Triticum monococcum* (einkorn wheat) subspecies are presented by alignment to *T. aestivum*, *T. urartu* and *H. vulgare* (Figure 1E). These resources contain between 118,000 to 140,000 transcripts each. The alignments to the selected reference genomes allows comparative analysis to be performed between the different resources, including ‘lift-over’ of features such as SIngle Nucleotide Polymorphisms (SNPs) from one species to another, described in (Fox et al., 2014).

## Analysis and visualisation

Every genome hosted by Ensembl Plants is subject to several automatic computational analysis, summarised in Table 3. Some of the key analysis and their resulting visualisations are described in more detail below.

**Table 3:** A list of the standard computational analyses that are routinely run over all genomes in Ensembl Plants.

| Pipeline name | Summary |
| --- | --- |
| Repeat feature annotation | Three repeat annotation tools are run, RepeatMasker, Dust and TRF.<br>RepeatMasker was run with repeat libraries from Repbase as well as Triticeae specific repeats from TREP.<br>http://ensemblgenomes.org/info/data/repeat_features |
| Non-coding RNA annotation | tRNAs and rRNAs are predicted using tRNAScan-SE (Lowe and Eddy, 1997) and RNAmmer (Lagesen et al., 2007), respectively. Other ncRNA types are predicted by alignment to Rfam models (Griffiths-Jones, 2005). |
|  | http://ensemblgenomes.org/info/data/ncrna |
| Feature density calculation | Feature density is calculated by chunking the genome into bins, and counting features of each type in each bin. |
| Annotation of external cross references | Database cross references are loaded from a predefined set of sources for each species, using either direct mappings or by sequence alignment.<br>http://ensemblgenomes.org/info/data/cross_references |
| Ontology annotation | In addition to database cross references, ontology annotations are imported from external sources (Ashburner et al., 2000; Cooper et al., 2013). Terms are additionally calculated using a standard pipeline based on domain annotation (Jones et al., 2014) and are projected between orthologues defined by gene tree analysis.<br>http://ensemblgenomes.org/info/data/cross_references |
| Protein feature annotation | Translations are run through InterProScan (Jones et al., 2014) to provide protein domain feature annotations.<br>http://ensemblgenomes.org/info/data/protein_features |
| Gene trees | The peptide comparative genomics (Compara) pipeline (Vilella et al., 2009) computes feature rich gene trees for every protein in Ensembl Plants.<br>http://ensemblgenomes.org/info/data/peptide_compara |
| Whole genome alignment | Whole genome alignments are provided for closely related pairs of species based on LastZ or translated BLAT results. Where appropriate, Ka/Ks and synteny calculations are included.<br>http://ensemblgenomes.org/info/data/whole_genome_alignment |
| Short read alignment | Short reads are automatically downloaded from the SRA by study accession and are aligned to a given reference by using BWA (Li and Durbin, 2009), GSNAP (Wu and Nacu, 2010), or STAR (Dobin et al., 2013), depending on the characteristics of the library. Alignments are stored in BAM or WIG format, and may be viewed as coverage or density tracks in the browser.<br>http://ensemblgenomes.org/info/data/short_read_mapping |
| Variation coding consequences | For those species with data for known variations, the coding consequences of those variations are computed for each protein-coding transcript (McLaren et al., 2010).<br>http://plants.ensembl.org/info/docs/tools/vep/index.html |

### Whole genome alignments

A total of 55 pairwise whole genome alignments are provided for the 20 monocot genomes in Ensembl Plants (Figure 4). Pairs include bread wheat against barley, *T. urartu* against *Ae. tauschii* and *Sorghum bicolor* against *Zea mays* and barley. Each genome was aligned to the *Oryza sativa* (Japonica) genome, allowing any pair of genomes to be indirectly compared via this reference. Additional comparisons include an all-against-all comparison of the 10 rice genomes, produced in collaboration with Gramene (Monaco et al., 2014).

**Figure 4:**
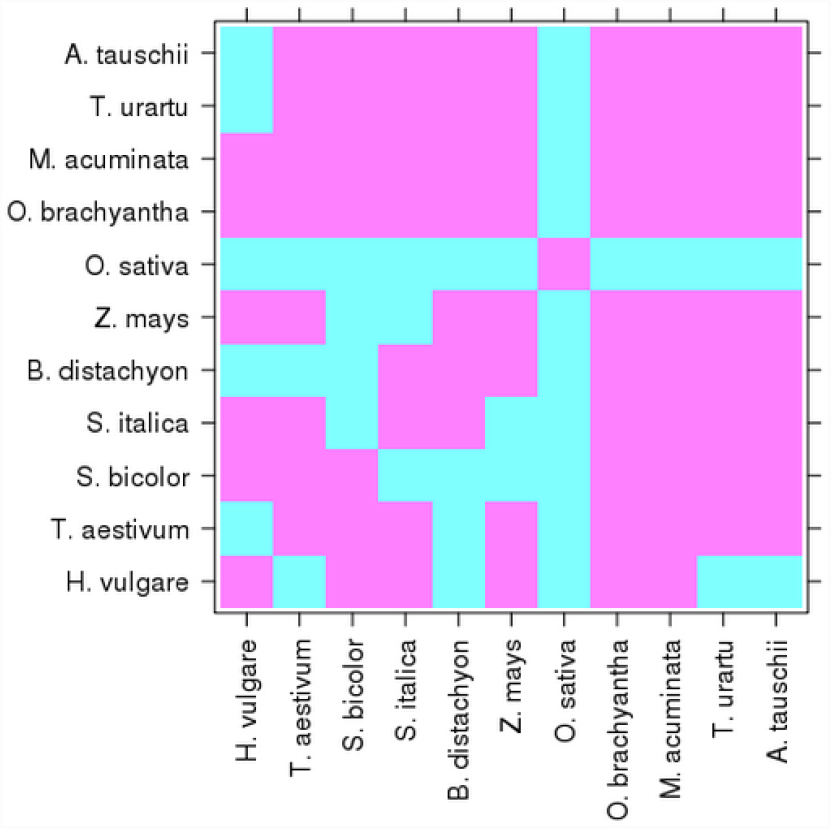

The matrix of whole genome alignments between pairs of monocot genomes in Ensembl Plants. Cyan indicates an alignment exists for the pair. Only one representative rice is shown, *O. sativa*, although each of the ten rice was aligned against each other (not shown).

In the first step of the whole genome alignment pipeline, two types of pairwise alignment may be used: either LastZ (Harris, 2007), for closely related species, or translated BLAT (Kent, 2002), typically for more distantly related species. After the initial alignment step, non-overlapping, collinear ‘chains’ of alignment blocks are identified, and the final step ‘nets’ together compatible chains to find the best overall alignment (Kent et al., 2003). When sequence similarity between the pair is sufficiently high, the ratio of the number of nonsynonymous substitutions per non-synonymous site (dN) to the number of synonymous substitutions per synonymous site (dS), which can be used as an indicator of selective pressure acting on a protein-coding gene (dN/dS) and synteny calculations are included and may be visualised in the genome browser.

Whole genome alignments are used to support the parallel visualisation of aligned genomic regions across multiple related species in the browser in the ‘Region Comparison View’ (Figure 5), allowing the inspection of conserved features and differences such as gene structure, copy number, polymorphism and repeat content between the genomes of multiple species.

**Figure 5:**
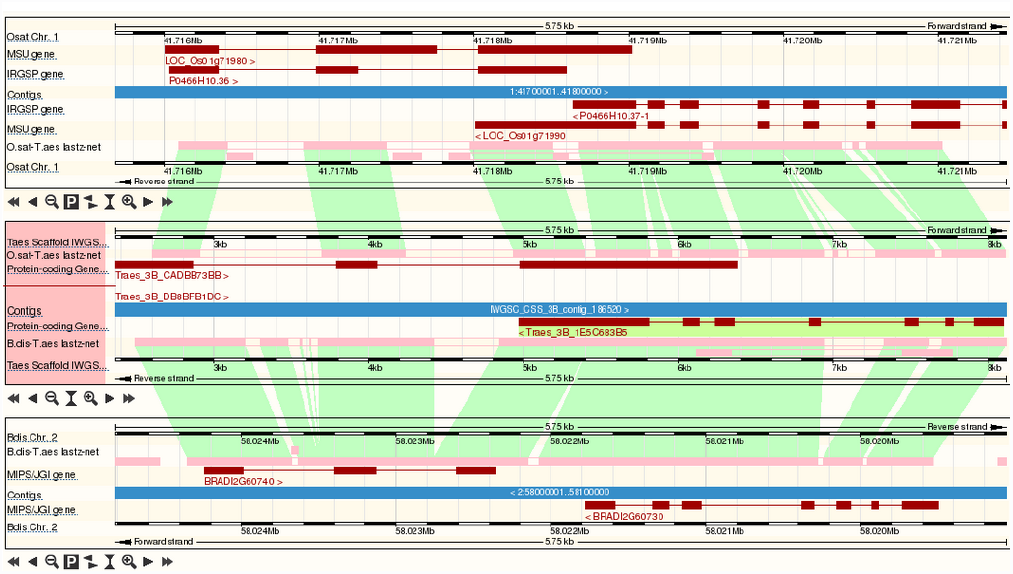

View of the whole genome alignment between wheat rice and brachypodium in Ensembl Plants. Pink bars and green blocks indicate aligned blocks between the rice and wheat (upper) and wheat and Brachypodium (lower) pairs of genomes. Transcripts are shown in red but the genomic features shown on each track are configurable.

Another potential use of whole genome alignments is the creation of functional ‘projection assemblies’ between a genome with a chromosome-scale assembly, such as brachypodum or barley, to one currently without, such as wheat. This task may be performed using BioMart in several steps. For example, two gene-based wheat markers may span a QTL of interest, but would likely not be located on the same contiguous assembly in the fragmented draft bread wheat assembly. However, one can retrieve the orthologues of these genes on the barley assembly where they are likely to belong to the same chromosomal region. In a second step, all the wheat orthologous of the barley genes in the region can be retrieved.

### Polyploidy

Building on the region comparison view for whole genome alignments, a recently developed ‘Polyploid View’ allows users to browse homoeologous genomic features on the wheat A, B and D component genomes in parallel (Figure 6). Alignments between contigs containing homoeologous gene family members, identified using gene trees (described below), can be directly visualised from the ‘Homoeologues’ page for each gene, and can be visualised with a single click. These alignment data are generated by comparing the three bread wheat genomes to each other using the same protocol as that used for the inter-species alignments. An additional filtering step retains only those alignments containing genes with inferred ‘orthology’ between the component genomes (defined as homoeologues, as described below). By using this definition, paralogous alignments between gene families are not show in the polyploid view.

**Figure 6:**
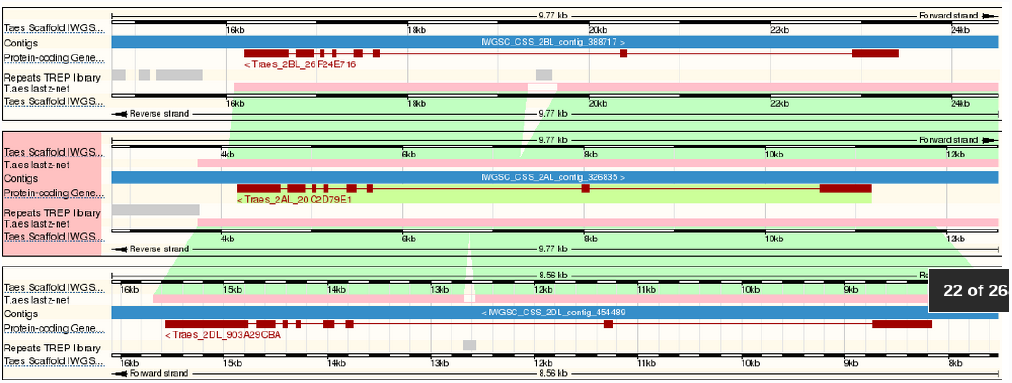

Polyploid view of the whole genome alignment within the bread wheat A, B and D component genomes. The image is defined as in Figure 5. An additional feature track shows repeats annotated in all three genomes.

### Gene trees

The Ensembl Gene Tree pipeline is used to calculate evolutionary relationships among members of protein families (Vilella et al., 2009). Clusters of similar sequences are first identified using BLAST+ (Camacho et al., 2009), then each cluster of proteins is aligned using M-Coffee (Wallace et al., 2006) or, when the cluster is very large, Mafft (Katoh et al., 2002). Finally, TreeBeST (Vilella et al., 2009) is used to produce a gene tree from each multiple alignment by reconciling the relationship between the sequences with the known evolutionary history to call gene duplication events. This final step allows the identification of orthologues, paralogues, and, in the case of polyploid genomes, homoeologues.

TreeBeST merges five input trees into one tree by minimizing number of duplications and losses into one consensus tree. This allows TreeBeST to take advantage of the fact that DNA-based trees often are more accurate for closely related parts of trees and protein-based trees for distant relationships, and that a group of algorithms may outperform others under certain scenarios. The algorithm simultaneously merges the five input trees into a consensus tree. The consensus topology contains clades not found in any of the input trees, where the clades chosen are those that minimize the number of duplications and losses inferred, and have the highest bootstrap support.

The resulting gene tree is an evolutionary history of a gene family, including identification of candidate gene duplication and speciation events, derived from the multiple sequence alignments. From the gene tree, one can identify true orthologues, paralogues and, in the case of polyploid genomes, homoeologues. Part of an example gene tree is shown in Figure 2, showing the inferred evolutionary history of a protein in the Sucrose-6F-phosphate phosphohydrolase family, including speciation and duplication events.

As a relatively large set of closely related genomes, the Poaceae (true grasses, of which the Triticeae are a subset) are particularly interesting. Using the gene tree analysis, we have placed 690,172 cereal genes into 39,216 groups of implied orthology. Inspite of the provisional nature of many of these genome assemblies, many of these orthologous groups are represented in every genome. A total of 7,203 groups (containing 260,004 genes) cover all 21 Poaceae genomes (counting the bread wheat A, B and D component genomes separately); 18,433 orthologous groups cover between 2 and 21 genomes with a single representative from each genome in the group; and 954 groups contain a single representative from all 21 genomes.

One of the benefits of extensive and accurate prediction of orthologs across plant species is the ability to project functional annotation between pairs of orthologous genes on the assumption that orthologues generally retain function between species (Altenhoff et al., 2012). Using this methodology we have projected manually curated Gene Ontology (GO) terms from *Oryza sativa* to the other monocots. Projected terms are tagged as “inferred from electronic annotation” (IEA) to prevent confusion with curated GO terms resulting from direct experimental evidence. The bulk of GO annotations for most species before projection are IEA assignments that come from Interpro2Go (see Table 2) that tend to be functionally broad. In contrast, projected terms can provide far more detailed annotation.

### Variation

The Ensembl Plants variation module is able to store variant loci and their known alleles, including SNPs, indels, and structural variations; the functional consequence of known variants on protein-coding genes; and individual genotypes, population frequencies, linkage and statistical associations with phenotypes. In the case of the polyploid bread wheat genome, heterozygosity, inter-varietal variants and inter-homoeologous variants are stored and visualised distinctly. A variety of views allow users to access this data (e.g. Figure 3) and variant-centric warehouses are produced using BioMart. In addition, the Variant Effect Predictor allows users to upload their own data and see the functional consequence of self-reported variants on protein-coding genes (McLaren et al., 2010).

## Future directions

The rapid pace of progress in the field of cereal genomics is driving the continued development of Ensembl Plants. Triticeae resources are a prioritised within the resource, and we aim to include new data sets rapidly after publication and data release. At the same time, the complex nature of these genomes is necessitating ongoing improvements to our analysis pipelines and user-interface.

Specific developments that are planned within the next few months include the release of improved genomic assemblies for barley and wheat. Although these assemblies will not be contiguous and ungapped, the use of additional genetic data will allow the approximate positioning and orientation of a larger number of genes within a chromosome-scale framework.

We expect to release an update to the barley genome assembly in release 24, due in October 2014, using a new marker set derived from population sequencing (POPSEQ) to anchor sequence contigs to chromosomal locations (Mascher et al., 2013). The new assembly will anchor an additional 346 Mb of sequence (411,526 contigs), containing 995 genes (5% of the total).

Similarly, we expect progress in the development of the wheat genome assembly towards full chromosome assemblies that makes use of both the recently released sequence of the 3B chromosome (Choulet et al., 2014) and integration of POPSEQ data across the whole genome (Poland et al., 2012; IWGSC, unpublished). Fuller assemblies will alleviate the computational burden of whole genome alignment, which is problematic when genomes are highly fragmented, allowing for the maintenance of a wider range of whole genome comparisons.

In addition to expanded variation data for bread wheat, by release 25, due in December 2014, we expect that the whole genome alignments between the A, B and D component genomes will be used to generate an extensive set of inter-varietal variations, and will be added to the existing variation data. Similar inter-species analysis based on the *T. monococcum* transcriptome will provide yet another source of wheat variation data.

In the longer term, we plan to extend the range and scope of RNA sequence alignments to the plant genomes hosted in Ensembl Plants by developing automatic methods to discover the relevant entries in the European Nucleotide Archive (Leinonen et al., 2011) based on their descriptive meta data. Matching datasets will be aligned automatically, and a new configuration interface will allow users to select studies to view against the relevant genomes based on matching search criteria against submission metadata. Work is also ongoing to integrate data and visualisation tools from the ArrayExpress (Rustici et al., 2013) and Atlas (Petryszak et al., 2014) resources into the browser, to allow expression data between tissues, time-series, or species to be viewed in a consistent way.

## Acknowledgements

Ensembl Plants is funded as part of the transPLANT project within the 7th Framework Programme of the European Union, contract number 283496. Databases are constructed in a direct collaboration with the Gramene resource, funded by the United States National Science Foundation award #1127112. D.M.B. wishes to thank Cristobal Uauy for the ‘projection assemblies’ use case.

